# GlycaNER: An Automated Glycan Structure Term Detection and Normalization Tool in Biomedical Literature

**DOI:** 10.64898/2026.11.13.766437

**Authors:** Shovan Bhowmik, Rene Ranzinger, Cyrus Chun Hong Au Yeung, Jeet Vora, Urnisha Bhuiyan, Michael Tiemeyer, Raja Mazumder, K. Vijay-Shanker

## Abstract

Glycans are complex carbohydrates covalently attached to biomolecules and play essential roles in protein folding, stability, intracellular trafficking, and molecular recognition. Glycan-related biological processes are vital to normal physiology, and their disruptions can contribute to various diseases. In the literature, glycans are described by their structures or recurring substructures, known as glycan structure terms (GSTs). We developed GlycaNER, a transformer-based entity recognition tool that detects GSTs and related glyco-terms in text, maps them to existing dictionary entries and suggests unmatched terms for expert curation. GlycaNER is developed with three objectives. First, it supports expansion of the Glycan Structure Dictionary (GSD), an existing but incomplete collection of experimentally verified GSTs with definitions, synonyms, and literature evidence, by identifying new entries and synonyms. Second, since textual mentions of glycan attachment described by GSTs often indicate glycosylation events, detecting these terms can aid extraction of glycosylation information from the literature. Third, because the literature links GSTs to biological functions and disease pathogenesis, GlycaNER can support tools to identify glycans’ impact on biological processes and disease. When evaluated on annotated abstracts, GlycaNER showed strong performance achieving an F1-score of 92.05%. To investigate scalability, we applied it to 10,000 PubMed abstracts, identifying over 50,000 glyco-term mentions, including approximately 14,000 unique terms. About half of all mentions were mapped to GSD entries, while over 8,000 unique mentions partially matched GSD entries. These findings highlight GlycaNER’s potential to reduce manual curation effort, expand the GSD, and enhance glycan resources for downstream biomedical research.

## Introduction

Glycosylation is a common type of post-translational modification in which complex carbohydrate moieties, known as Glycans, are covalently attached to specific amino acid residues within a protein (Stowell et al. 2015). Glycans play important roles in protein folding, stability, intracellular trafficking, and molecular recognition (Moremen et al. 2012). Through these diverse functions, glycans contribute to the maintenance of normal cellular and physiological processes (Varki 2017). Alterations in their synthesis and pathways can cause the initiation, progression, and manifestation of a broad range of diseases including congenial disorders (Ng and Freeze 2018; Reily et al. 2019).

Large numbers of glycans have been identified experimentally, exhibiting structural diversity with complex branching patterns and multiple glycosidic linkages (Cummings 2009; Moremen et al. 2012). Glycans are often described in a partial manner in the literature as motifs, types, subtypes, branching, terminal structures, or recurring sub-structures. These descriptions are collectively referred to as Glycan Structure Terms (GSTs) in a previous study (Vora et al. 2023). In this work, the Glycan Structure Dictionary (GSD) was developed to organize experimentally verified glycan-related terms, including GSTs and glycoconjugates (GCTs) along with their cross-references, synonyms, and supporting publications. GCTs are molecules in which glycans are covalently linked to non-carbohydrate components, such as proteins or lipids (Shivatare et al. 2022). As GCTs typically provide information about the attached glycan or its structure, these terms were also included in the GSD. The terms included in the GSD are primarily collected from the scientific literature through a semi-automated text-mining process.

Although GSD provides a curated collection of glyco-terms, its coverage is not comprehensive. New terms and textual variants continue to emerge in scientific literature. Automated recognition of such terms from text is therefore important for further expansion and maintenance of GSD. The GSD development (Vora et al. 2023) is a primary motivation for developing GlycaNER, addressing the need for recognizing and extracting a broader range of glyco-terms and their textual variants from the text with the goal of further expansion of GSD. Additionally, glycosylation events are often indicated in the literature by mentions of attached glycans described using GSTs (Kolarich et al. 2006; Vora et al. 2023). For instance, in PMID 16622833, *“…exclusively diantennary structures were present on site N271 in each of the isoforms analyzed…”*; the GST *“diantennary structures”* describes an N-linked glycan structure covalently attached to the *“N271”* residue, indicating a glycosylation event. Thus, GlycaNER can serve a role in glycosylation information extraction from the literature. Moreover, glycan properties such as branching patterns, terminal features, core structures, represented by GSTs can influence biological functions, protein interactions and disease pathogenesis and may serve as clinical biomarkers (Dennis et al. 2009; Pinho and Reis 2015; Varki 2017; Reily et al. 2019). Thus, identifying GSTs in publication can facilitate such research by linking glycans to relevant biological information.

The importance of glycans has led to the development of several glycoinformatics resources, including CarbBank (Doubet et al. 1989), GLYCOSCIENCES.de (Lütteke et al. 2006), and CFG (Raman et al. 2006), which established early glycan databases. GlyCosmos (Yamada et al. 2020) created a unified portal linking glycans to pathways and diseases. Other resources include GlyConnect (Alocci et al. 2019) for collecting glycoproteomics data and GlyGen (York et al. 2020) for integrating glycoscience knowledge.

The significance of glycan structural terms has given rise to the development and characterization of several resources such as GlyTouCan (Fujita et al. 2021) as a structure-focused repository for glycan structures, providing unique accession numbers for consistent referencing; GlycoMotif (GlycoMotif 2025) as a resource for common motifs and substructures found inside larger glycans, and GlycoEpitope (Okuda et al. 2016) covering glycan epitopes.

In this work, we developed GlycaNER, a transformer-based named entity recognition (NER) tool that first uses a fine-tuned BioBERT model (Lee et al. 2020) to detect glyco-terms in text. BioBERT (Lee et al. 2020) and BioMedBERT (Gu et al. 2021), which are adaptations of the transformer language model BERT (Devlin et al. 2019) to the biomedical domain, are widely used frameworks for developing biomedical NER systems. Once detected, each recognized entity is assigned to its corresponding entity type. In this research, we consider five distinct types of glyco-terms: GSTs; GCTs; glycan sequences; compositions; and simple or modified sugars (definition of these terms along with examples are provided in Supplementary Section S1). Although only GSTs and GCTs are represented in the GSD, the recognition of remaining types can provide useful information for extracting glycosylation events from the literature.

GlycaNER also determines whether each detected entity is structurally informative or non-informative. Non-informative terms usually refer to glycan classes or structures without specifying structural information. They are frequently found in the literature and include terms such as *“glycan”, “carbohydrate” and “determinant”.* We will call such terms *generic*. GlycaNER further checks for matches with existing GSD entries. If a match is found, GlycaNER returns the corresponding GSD entry id; otherwise, the entity is considered a candidate for potential inclusion in the GSD.

Previous work on extracting glyco-terms directly from text remains limited. More recently, the GlyCosmos group used PubAnnotation to screen more than 15,000 abstracts for glycan motifs and epitopes, together with six other biological entity types including phenotypes, anatomy and diseases. They integrated the resulting annotations into the GlyCosmos portal (Kim et al. 2026). Their approach used predefined dictionaries of glycan motifs and epitope names to identify these entities in text. In contrast, GlycaNER is designed to automatically detect a diverse range of glyco-terms, including glycan motifs and epitopes, from text without relying on a predefined vocabulary.

In this research, we conducted two studies to evaluate GlycaNER. First, we assessed its accuracy in detecting glyco-terms and correctly identifying their entity types. The results demonstrated strong performance, with GlycaNER achieving an overall F1-score of 92.05%. Second, we examined the large-scale application of GlycaNER by applying it to a corpus of 10,000 abstracts. This study explored its ability to capture a wide range of glyco-terms, including those not currently represented in the GSD. It also allowed us to assess the potential of GlycaNER to identify candidate terms for future GSD expansion. GlycaNER identified 51,874 glyco-term mentions, representing 14,023 unique terms, and 24,217 mentions were successfully normalized to existing GSD entries.

The Results section below presents the detailed evaluation and large-scale application of GlycaNER. We then discuss the findings and implications of our tool in the Discussion section, followed by the Methods section that describes the GlycaNER pipeline and its individual components.

## Results

### Study 1: Performance Evaluation of GlycaNER

Our first study evaluates GlycaNER’s ability to detect glyco-terms in unseen text and accurately classify the detected entities into their corresponding types. Our evaluation was conducted on a set of 100 abstracts that are included as literature evidence for GSD entries. Although each abstract served as publication evidence for a single GSD entry, experts annotated all other relevant terms found in these abstracts. These 100 abstracts contained 1,007 glyco-terms across the five entity types detected by GlycaNER: 391 GSTs, 368 GCTs, 144 simple and modified sugars, 96 glycan sequences, and 8 glycan compositions. These 100 abstracts used for evaluation are distinct from those used to train our BERT model.

Following standard practice for biomedical NER systems (Leaman and Gonzalez 2008; Doğan et al. 2014; Lee et al. 2020), we report precision, recall, and F1-score computed from true positives (TP), false positives (FP), and false negatives (FN). A GlycaNER-detected string was counted as a TP if (i) it matched an annotated string exactly and (ii) GlycaNER assigned the same type to the string as per the annotations. The first criterion is stricter than the partial-match criterion commonly used in NER evaluations, which considers a detected string correct if it overlaps with an annotated string. Thus, a detected string that only matches partially or does not match an annotated string is considered an FP. Similarly, an FN arises when GlycaNER fails to detect an annotated term *exactly* or assigns a type different from the annotation. If GlycaNER only partially detects an annotated term or the assigned type does not match the annotated type, we count it as both an FP and an FN.

As mentioned in the “Introduction” section, generic terms contain limited information. They occur frequently in text and are easily detectable. Therefore, we exclude all terms identified as generic by GlycaNER from the evaluation.

From these counts, precision, recall, and F1-score were computed as follows:

*Precision = TP / (TP + FP); Recall = TP / (TP + FN); F1-Score = 2 × (Precision × Recall) / (Precision + Recall)*

We evaluated the performance for each type of glyco-term individually detected by GlycaNER. We also calculated the overall performance by measuring micro-averages, where the TP, FP, and FN count from all types were combined before calculating Precision, Recall, and F1-score, instead of simply averaging the scores across types. Table I shows the performance of GlycaNER on the evaluation set and will be discussed later.

**Table I.** Overall performance of GlycaNER in detecting multiple types of non-generic glyco-terms in 100 abstracts.

| **Type** | **Precision (%)** | **Recall (%)** | **F1-Score (%)** |
|---|---|---|---|
| GST | 91.35 | 91.82 | 91.58 |
| GCT | 98.83 | 91.85 | 95.21 |
| Sequences | 84.27 | 78.13 | 81.08 |
| Composition | 100 | 100 | 100 |
| Simple and Modified Sugars | 100.00 | 85.42 | 92.13 |
| **Overall** | **94.55** | **89.67** | **92.05** |

### Study 2: Large-Scale Extraction from PubMed

In addition to evaluating GlycaNER’s accuracy, we also investigate whether we can run our tool at a larger scale to recognize diverse glyco-terms from the literature. To obtain this large set of PubMed abstracts, we first constructed a PubMed query as discussed below.

### Query Construction

To design a PubMed query likely to return a large set of abstracts that contains a wide range of terms, we included all sub-components of glycan structural terms as described in the “Decomposition” section. By including sub-components rather than full GSD entries, the query can retrieve abstracts containing glycan-related terms beyond those already present in GSD. Structural terms that are also common in other scientific contexts, such as “core 1”, “type 2”, and “hybrid”, were included only in the query provided they co-occur with another glycan structural component or a complete GSD entry within the title or abstract. All the sub-components that we collected are connected by “OR”. The complete PubMed query, along with the inclusion and exclusion criteria, is provided in Supplementary Section S2.

### GST Coverage and Distribution

Our constructed PubMed query returned 373,096 published abstracts. From these records, we randomly selected 10,000 abstracts for this study. Our tool identified at least one glycan-related term in all but 420 of the 10,000 sampled abstracts, extracting a total of 51,874 terms after excluding 10,828 generic terms. GST was the most frequently detected category, followed by GCT and other glycan-related types.

To assess whether GlycaNER could capture additional synonyms and novel terms beyond the existing terms already included in the GSD, the extracted GSTs and GCTs were standardized against GSD. Of these, 24,217 mentions matched a dictionary entry exactly, while 15,444 showed a partial match. The exact matches were normalized to 148 unique GSD entries, whereas the partial matches corresponded to 8,207 unique terms beyond the curated dictionary. Table II summarizes the distribution of five types of glyco-terms, showing the total number of mentions extracted by GlycaNER from the literature and the number of unique terms identified among these mentions.

**Table II.** Distribution of non-generic glycan-related terms detected by GlycaNER across 10,000 abstracts sampled from the results of the constructed PubMed query.

| **Type** | **Total Mentions** | **Unique Terms** |
|---|---|---|
| GST | 26,617 | 9,319 |
| GCT | 12,751 | 1,804 |
| Sequences | 1,870 | 1,507 |
| Composition | 167 | 70 |
| Simple and Modified Sugars | 10,469 | 1,323 |
| **Total** | **51,874** | **14,023** |

## Discussion

### Study 1

As shown in Table I, GlycaNER demonstrates strong performance in detecting glyco-terms, achieving nearly 95% precision and 90% recall. The recognition of the two major types of glyco-terms in the GSD (GSTs and GCTs), as well as the recognition of simple and modified sugars, achieved F1-scores exceeding 90%. Composition detection achieved an F1-score of 100%.

For glycan sequences, some annotated strings were very long. The BERT model could not recognize these strings properly and mostly identified a portion of the complete sequence. Because the shorter predicted strings did not match the actual annotated entities, they were counted as FPs, while the complete annotated sequence remained undetected and were counted as FNs. This type of partial detection was the primary reason for the lower precision and recall observed for glycan sequences compared to the other glyco-term types.

Although no FPs were observed for sugar entity detection (100% precision), one source of FN error was the handling of acronyms specified within parentheses. Consider the following phrase, “*…3’-galactosyl-lactose (beta3’-GL), 4’-galactosyl-lactose (beta4’-GL), and 6’-galactosyl-lactose (beta6’-GL)…”* from PMID 25063655. In this case, the BERT model correctly identified the full terms (“*3’-galactosyl-lactose”, “4’-galactosyl-lactose” and “6’-galactosyl-lactose”*) but failed to recognize the corresponding acronyms: “*beta3’-GL”*, “*beta4’-GL”*, and “*beta6’-GL”*. We did not use any acronym detectors in this work, and believe that its usage could have eliminated many of these FNs. In the future, we plan to incorporate acronym detection to reduce such errors which could also appear in other types.

### Study 2

As shown in Table II, GlycaNER extracted over 50,000 non-generic terms from 10,000 abstracts. As expected, GSTs were the most frequently detected type, with more than 25,000 mentions. Among these, more than 9,000 terms were unique. Out of these unique GSTs, some examples from the top ten most frequently occurring GSTs in this set of abstracts were *“Heparan Sulfate”, “Sialyl Lewis X”, “Sialyl-Tn Antigen”, “biantennary N-glycans”, and “SSEA-4”*. GCT was the next most common type, with more than 12,000 mentions, of which 1,804 were unique. Few frequently extracted GCTs include *”GM1”, “Asialo-GM1”, “Neolactotetraosylceramide”, “Sulfatide”, and “9-O-Acetyl-GD2”*. All these ten terms are found in the current GSD.

For the other entity types, Table II shows that most detected sequences were unique. We also identified around 1,300 unique simple and modified sugars. In contrast, composition mentions were much less frequent than the other entity types. GlycaNER identified 70 unique compositions in the large-scale analysis.

Of all detected terms, more than 24,000 mentions were normalized to GSD entries or their synonyms. Of interest to us, more than 15,000 terms were partial matches to GSD entries. Among these partial matches, more than 8,000 were unique. Inspection of the partial matches revealed several GSTs and GCTs which could potentially be included as new entries in the GSD. Some examples are shown below:

- ***“4-O-Acetyl-Sialic Acid”:*** partially matches with an existing GSD entry “9-O-Acetyl Sialic Acid”. Given the similarity with the existing GSD term, we can assume that 4-O-Acetyl Sialic Acid is a reasonable potential candidate as a sibling term in the GSD.
- ***“3’-Sulfo-Lewis C”:*** although there are Lewis X, Lewis A and their variants in the dictionary, there are no entries involving Lewis C or any of its structural variants. Again, we consider this mention as a potential candidate for inclusion in the GSD.
- ***”2,4-triantennary oligosaccharides”:*** partially matched the existing GSD entry *”Triantennary N-glycans”*. The detected term contains additional linkage information *“2,4”* absent in the existing entry and could therefore be considered a separate or a replacement of the existing GSD entry. Another such example is the mention *”Mono-sulfated isoglobotetraosylceramide”* that matches partially with an existing GSD entry *”Isoglobotetraosylceramide”* with additional information about the sulfation of the *“isoglobotetraosylceramide”* glycosphingolipid.
- **“*difucosyl type 1 chain A”:*** This represents a different situation where the detected entity does not match any existing GSD entry. In this case, although it might not be included as a new GSD entry, it may qualify to be added as a synonym for an entry *”Blood Group A (Type 1) - Lewis B”* which already exists in the GSD.

These examples represent only a small sample of the thousands of unique partial GSTs and GCTs detected by GlycaNER. We plan to send all the partial matches to the experts for their verification and inclusion in the GSD. Thus, Study 2 demonstrates GlycaNER’s potential to address its main motivations: assist in developing a more comprehensive GSD by identifying new glyco-terms and synonyms of the existing terms; facilitating the identification of glycan attachments for glycosylation information extraction; and advancing research linking glycans with biological functions, processes, and/or diseases.

## Materials and methods

The GlycaNER pipeline is shown in Figure 1.

**Figure 1.**
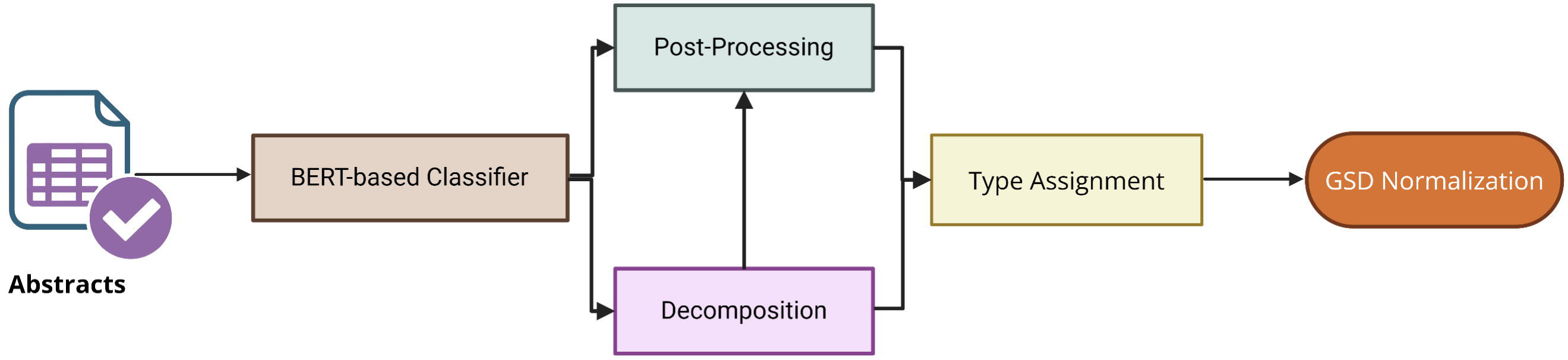
Overview of the GlycaNER application pipeline. PubMed abstracts are processed by a BERT-based classifier to identify candidate glyco-terms, which are refined through post-processing and assigned types. Decomposition into glycan structural components supports both post-processing and subsequent normalization of the detected terms against GSD entries and their synonyms. **Alt text:** Flowchart showing the GlycaNER pipeline, in which PubMed abstracts are processed by a BERT-based classifier to identify candidate glyco-terms. The detected terms undergo post-processing and type assignment. Glycan structural components are then extracted and used for post-processing and normalization against GSD entries and their synonyms.

In the following, we describe each step, starting with its purpose and motivations.

### Fine-Tuned BERT Model

The processing starts with the application of a transformer-based BERT encoder, a central part of the pipeline, which processes sentences from PubMed abstracts and identifies candidate strings.

In GlycaNER, we use BioBERT (Lee et al. 2020), a BERT-based language model specialized for the Biomedical domain. This language model is trained to identify glycan-related terms using the BIO tagging scheme, which is widely used for NER tasks (Zhang and Chen 2022). The annotated dataset used for training the classification model labels each word/token with *B (beginning), I (inside), or O (outside)* labels. For example, in the word sequence below, “*6-sulfated sialyl Lewis x is a major capping group of GlyCAM-1”, “6-sulfated sialyl Lewis x”* is a glycan-related string. Here, the token *“6-sulfated”* is labeled as *“B”*, *“sialyl”, “Lewis” and “x”* are labeled as *“I”* and all other tokens in the sequence are labeled as *“O”*. Thus, the labeling indicates that “*6-sulfated sialyl Lewis x*” should be identified as a glyco-term, whereas the remaining tokens, labeled as *O*, are not part of a named entity.

### Training the BERT model

The training dataset for the BioBERT language model was a corpus annotated by domain experts. To create the training dataset, we collected 601 abstracts randomly chosen from those included as publication evidence for GSD entries. This training corpus comprising 601 abstracts contains 5,200 sentences that have 4,725 annotated terms.

To identify a choice of hyperparameters for the model, we used an 80-20 split of the entire training dataset for training and development. After testing different sets of hyperparameters, the following set of hyperparameters was picked: learning rate: 5e − 5, batch size: 16, optimizer: AdamW, L2 regularization: 2e − 5, epochs: 5, loss function: Cross-Entropy.

### Decomposition

To facilitate the normalization process, which involves matching a detected string with strings in GSD, we decompose both the detected string and the strings in GSD into meaningful component units. The decomposition process uses a curated list of structural components belonging to several glycan structural classes and sub-classes, including glycan structural descriptors, named glycan motifs, sugar elements, chemical substituents and glycoconjugates (See “Glycan Structural Component” subsection below). When decomposing a string into meaningful units, we use the curated structural components to identify the longest matching component. For example, the GSD entry “2,6-branched O-mannose” is split into “*2,6”* (Positional Descriptor), *“branched”* (Topological Descriptor), *“O”* (Linkage Descriptor), and *“mannose”* (Monosaccharide). This decomposition facilitates normalization by allowing related forms of the same components to be matched. For instance, *“mannose”* in the above-mentioned GSD entry can be matched with its adjectival form *“mannosyl”* in a GlycaNER detected string *“2,6-branched structure in brain O-mannosyl glycan”* (PMID: 14617637), as both belong to the same sugar class and can be standardized to a common form (See “Normalization” section).

Similarly, another string *“alpha-(2,3)-sialylated core-fucosylated complex type bi-antennary”* (PMID: 18601234) is decomposed into *“alpha”* (Linkage Descriptor), *“2,3”* (Positional Descriptor), *“sialylated”* (Modification Pattern), *“core-fucosylated”* (Core Structure Descriptor), *“complex type”* (Glycan Compositional Descriptor), *“bi”* (Numeric and Multiplicity Descriptor: Prefix), and *“antennary”* (Topological Descriptor).

Note that, all the descriptors are associated with the structural components that they modify. Thereby, preserving the relationships between components during the decomposition. For instance, in *“3′-Sialyl-6′-Sulfo Lewis X”*, the positional descriptor *“3′”* is associated with *“Sialyl”*, while *“6′”* is associated with *“Sulfo”*. Maintaining these associations ensures that the positional information is correctly linked to the corresponding structural modification during normalization.

While decomposition is central to the normalization process, it also assists in the post-processing of BERT-recognized entities by generating multiple valid glyco-terms from a single-entry containing terms connected by conjunctions or parentheses (discussed in “Post-Processing” section).

### Glycan Structural Components

To provide a basis to decide how to decompose a predicted glyco-term, we developed a curated set of *Glycan Structural Components.* Based on our examination of GSD entries, we curated these components into several glycan structural classes and subclasses, including glycan structural descriptors, named glycan motifs, sugar elements, chemical substituents, and glycoconjugates. These classes and their associated components define the splitting of tokens used in the decomposition process described above.

Some of these classes are also split into multiple sub-classes. The components of these classes include dictionary-based named entities as well as regular expression-based patterns used to capture glycan structural or recurrent sub-structural units.

The complete list of glycan structural classes and their components, used to identify parts within the detected glyco-strings and to split GSD entries for matching, is numerous and provided in Supplementary Section S3.

### Post-Processing

This step is completed after candidate strings are obtained from the BERT classifier and have undergone decomposition. This step serves two purposes, filtering some terms selected by the BERT model and refining the selected strings in some cases.

### Filtering non-glyco terms

Although the BERT model is trained to predict glyco-terms, some predicted strings may occur as part of other biological entities such as proteins and complexes, cell lines, antibodies, and phenotypes. As is common in NER, we assume that parts of longer entity names should not be detected separately. For example, in PMID 19267921, the BERT model identifies *”core 2”* as a GST although it appears as part of another entity name *”…the core 2 beta-1,6-N-acetylglucosaminyltransferase…”. “core 2”* often appears in the training set, so its prediction as a GST is not surprising. However, we aim to filter out these cases where a predicted term occurs as part of a longer name that does not correspond to one of the required glyco-term types in this work.

To filter such cases, we extract the base noun phrase from the corresponding BERT prediction. The base noun phrase represents the smallest noun phrase that includes the main noun. The main noun (also called the head noun) is typically the last word in a base noun phrase in English and usually carries information about its type. Since the head noun in the above example ends with a suffix indicating a protein, we omit the selection of *“core 2”* from the final output.

We have created a list of nouns that represent non-glyco entities. If the head noun of a base noun phrase belongs to one of our listed nouns, then we discard the candidate phrase detected by the BERT model. We use “spaCy” (Srinivasa-Desikan 2018), an open-source NLP library, to detect the base noun phrases.

### Refining BERT predictions

The second part of post-processing refines the BERT-predicted spans into correct GSTs that can be matched against the GSD entries. This step handles conjunctions and parenthetical expressions found in the BERT-obtained candidates.

#### Conjunction Resolution

Although BERT-recognized strings typically correspond to a single entity, they may sometimes contain multiple terms connected by conjunctions. For example, in PMID: 10460829, our BERT-predicted term is *“Forssman penta- and tetraglycosylceramide”*, which corresponds to two entities: *”Forssman pentaglycosylceramide”* and *”Forssman tetraglycosylceramide”*. Decomposing terms based on glycan structural components helps resolve terms connected by conjunctions. In the above scenario, *“tetraglycosylceramide”* is decomposed into “tetra,” “glycosyl,” and “ceramide.” This decomposition allows us to identify “penta” and “tetra” as components belonging to the same structural category and that they should occupy similar positions in the newly generated entities. Our method then distributes the remaining tokens across both entities, resulting in the two terms mentioned above. Similarly, the BERT-detected string *“3’-sialyl, 6’-sulfo, 6-sulfo, or 2’-sulfo LacNAc”* (PMID: 1429720) was resolved into four distinct entities: *“3’-sialyl LacNAc,” “6’-sulfo LacNAc,” “6-sulfo LacNAc,”* and *“2’-sulfo LacNAc.”*

#### Parentheses Resolution

Some BERT-predicted candidate strings contain parenthetical expressions. The part within the parentheses shouldn’t be considered part of the main glycan-related term but might be used to create an alternate term.

For example, the BERT-predicted string *“isoglobotetraosylceramide (iGb4Cer) IV3-sulfate”* in PMID: 1396705, is resolved into two distinct entities: *“isoglobotetraosylceramide IV3-sulfate”* and *“iGb4Cer IV3-sulfate.”* Here, the term outside the parentheses and its acronym within the parentheses are treated as alternative components. Then, “IV3-sulfate” is propagated to produce two terms.

On the other hand, parenthetical expressions containing quantitative values are removed because they are not part of the glyco-term. Thus, *in PMID:* 1429720, instead of *“3’-sialyl LacNAc (12.4%)”*, our glyco-term is “*3’-sialyl LacNAc”*. However, in PMID: 8486686, *“9(7)-O-acetyl GD3”* is retained as a single entity. Here, the parenthetical value denotes an alternative position of the O-acetyl group and is integral to the entire entity. Since our treatment of parentheses is to address acronyms, when an expression within parentheses contains numbers, we do not believe that they could be acronyms. Hence, we do not make any changes to the given mention.

### Type Assignment

After determining the final glyco-term, we then classify them into different types: GSTs, GCTs, glycan sequences and compositions, and sugars and their modified forms. Our assignment of the above-mentioned types involves a combination of rule-based heuristics, regular expression patterns, and dictionary-based matching. GSTs, GCTs, sugars, and generic terms are identified using the glycan structural classes and headwords listed in Supplementary Section S3. Sequences and compositions are detected by separate heuristics as they are not written as English phrases rather follow their own patterns. A sequence combines a residue abbreviation such as Gal, GlcNAc, or Neu5Ac with a linkage pattern, an arrow, or an anomeric descriptor, while a composition consists of two or more residue names each carrying a numeric count, as in Hex5HexNAc4Fuc1.

We additionally classify terms as *“Generic”* when they lack any specific structural features and are commonly found in the glyco literature. Although we did not include generic terms in either of our studies, we still detect them because they are glyco-related terms and may indicate glycosylation events.

### Normalization

After assigning types to each glyco-term, we normalize the terms against existing GSD entries. For each detected glyco-string, we identify glyco-relevant tokens and decompose the string into its component parts (See the “Decomposition” section for more details). GSD entries are similarly decomposed in advance. Headwords that do not contribute to the normalization process are excluded from the matching. The remaining components are used to retrieve candidate GSD entries through an inverted index, a lookup data structure that maps each component to all the GSD entries that contain it (Søgaard et al. 2015). To facilitate this matching, each component is further standardized to a common term. The standardized component parts of the detected string are then compared with those of the GSD entries. If the complete set of relevant components matches exactly, then the corresponding matched GSD entry’s ID is linked to the detected term. Terms that partially match or share no common common components with existing GSD entries can be forwarded to experts for manual curation. Figure 2 illustrates this normalization process.

**Figure 2.**
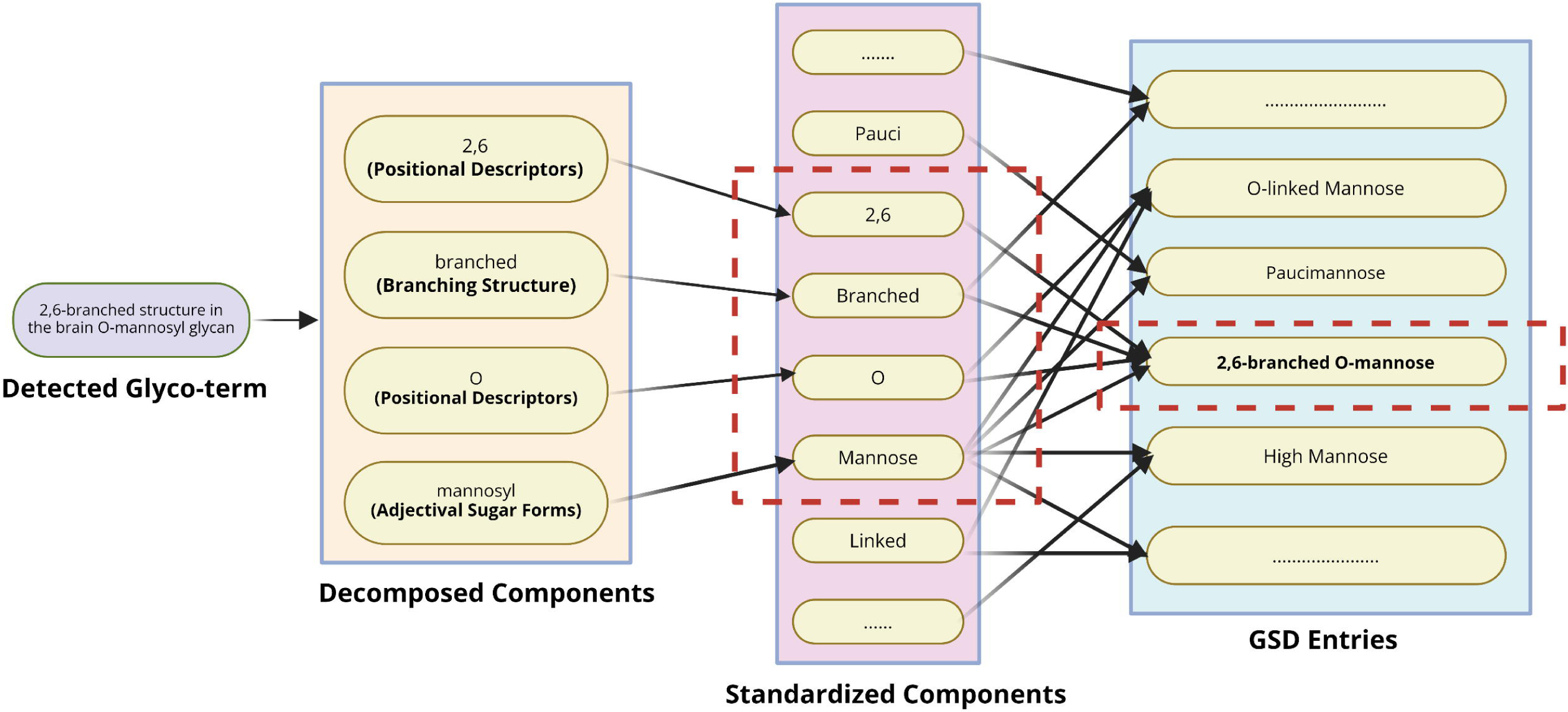
Inverted index-based normalization of a GlycaNER-detected glyco-term to a GSD entry. The detected glyco-string *“2,6-branched structure in brain O-mannosyl glycan”* is decomposed into structural components, which are standardized to common forms and matched against pre-computed components of GSD entries using an inverted index. The shared components *“2,6,” “Branched,” “O,”* and *“Mannose”* identify *“2,6-branched O-mannose”* (GSD000001) as the corresponding normalized GSD entry. **Alt text:** Diagram showing how the GlycaNER-detected term “2,6-branched structure in brain O-mannosyl glycan” is decomposed into standardized structural components and matched to pre-computed GSD entry components using an inverted index. The shared components “2,6,” “Branched,” “O,” and “Mannose” identify “2,6-branched O-mannose” (GSD000001) as the normalized entry.

We now illustrate the normalization process by describing how we normalized the detected string “*2,6-branched structure in brain O-mannosyl glycan”* to the corresponding GSD main entry *“2,6-branched O-mannose”* with accession ID GSD000001 (Vora et al. 2023). The normalization process begins by decomposing the mentioned glyco-string into its relevant components: “*2,6”, “branched”, “O”, “mannosyl”,* while non-contributory tokens such as *“structure”, “brain” and “glycan”* are excluded from matching. The corresponding GSD entry, which was decomposed in advance, contains “*2,6”, “branched”,* “O” and “mannose”. The components are then standardized, such that *“mannosyl”* and *“mannose”* are both represented as *“Mannose”.* The resulting components *“2,6,” “Branched,” “O,”* and *“Mannose”* retrieve several candidate GSD entries through the inverted index. Among these candidates, *“2,6-branched O-mannose”* contains the complete set of matching components and is therefore selected as the normalized term, with accession ID GSD000001.

The example above shows how detected glyco-terms can be automatically linked to existing GSD entries through component-based normalization. As mentioned in the “Results” section, at scale, approximately half of the detected mentions were successfully normalized to existing GSD entries, while over 8,000 unique mentions received partial matches, highlighting their potential for future inclusion in the GSD.

## Acknowledgments

We thank all current and past GlyGen developers and collaborators for the continuous development of GlyGen knowledgebase and software tools, which have greatly facilitated this work.

## Data availability statement

All original data and source code generated and used in this study are available at: https://github.com/glygener/GlycaNER

## Supplementary Data

**S1: Summary of glyco-term Types**

**S2: Query Construction for large-scale extraction study S3: List of Glycan Structural Components**

