## Supplementary Data for "GlycaNER: An Automated Glycan Structure Term Detection and Normalization Tool in Biomedical Literature"

**Running Head:** GlycaNER: An Automated GST Detection and Normalization Tool

### Supplementary Data

#### S1: Summary of glyco-term Types

Table S1.1 summarizes five non-generic glyco-terms that GlycaNER can extract from the literature.

**Table S1.1.** *Glyco-term types recognized by GlycaNER*

| Type | Definition | Example |
|---|---|---|
| **Glycan Structure Term (GST)** | Partial description of a glycan by its motif, type, subtype, branching, terminal features, core and recurring sub-structures. | Sialyl Lewis X; Fucosylated bisialo biantennary; |
| **Glycoconjugates (GCT)** | Molecules in which a glycan is covalently attached to a non-carbohydrate moiety. | GM1; Globopentaosyl Ceramide; |
| **Sequences** | Explicit residue-by-residue notation with anomeric and linkage information. | “NeuAc alpha 2-6Gal beta 1-4GlcNAc”; “Gal beta 1-->4GlcNAc beta 1-->6Gal”; |
| **Compositions** | Two or more monosaccharide, modified sugar, or sugar-derivative residues associated with explicit numeric counts. | GalNAc(2)GlcNAc(2)Man(3) Fuc(1) |
| **Simple and Modified Sugars** | Simplest forms of carbohydrates (monosaccharides) along with disaccharides that are formed by glycosidic linkages of two monosaccharides. Chemical modifications of these sugars are also part of it. | Fucose; 3-O-Sulfated Gal; UDP-GlcNAc; |

#### S2: Query Construction for large-scale extraction study

In addition to evaluating GlycaNER on an annotated set of abstracts, we conducted a second study to test the system's expandability on a large set of abstracts. This study aimed to extract a wide variety of glyco-terms from the literature. To identify glycan-related articles from millions of PubMed records, we constructed a query designed to capture a broad range of literature related to glycans and their structural components.

To construct the query, we used almost all the structural component terms included in our normalization process (see Supplementary Section S3). However, generic headwords that do not carry structural meaning on their own were excluded (e.g., antigen, ceramide, and structures). Named glycan entities, such as Forssman antigen and T antigen, were included as complete phrases in the query.

Some components were included to broaden the search but required additional constraints. These included terms such as core 1 to core 4, hybrid, complex, branch, sulfate, dimeric, and bisecting. These terms are not specific to glycans and can also appear in other chemical descriptions. Therefore, we required them to appear with at least one other glycan-related term.

Once the query was constructed, we used it to search PubMed titles and abstracts. We applied no restrictions on publication year, language, or article type. The search returned 373,096 records published between 1913 and 2026. From these records, we randomly selected 10,000 abstracts for large-scale glyco-term extraction using GlycaNER.

#### S3: List of Glycan Structural Components

GSTs are typically combinations of structural components used to describe glycans. We curated a list of glycan structural classes and the components belonging to each.

While we constructed the following component classes by analyzing a range of glyco-terms, we recognize that some classes may not yet be comprehensive. However, the class structure itself is fixed. Therefore, any future work would focus on expanding the content of the existing classes by adding missing components, rather than introducing new classes or modifying the current classification scheme.

##### Class A: Glycan Structural Descriptors

Recurrent glycan nomenclature elements and structural descriptors commonly used, including multiplicity indicators, linkage descriptors, antigenic determinants, and glycan series identifiers.

***Topological descriptors***

Topological roles that describe where a residue or motif is positioned within a glycan structure.

antenna | antennae | antennary | antennaries

branched | branching

terminal

termini

bisecting

***Numeric and multiplicity descriptors (Prefix)***

(bi | di | dim | tri | tria | tetra | pent | penta | hexa | hex | hept | hepta | octa | nona |

mono | poly | oligo) (meric)?

***Positional descriptors (Prefix)***

\d-\d-?

\d-?

***Linkage descriptors (Prefix)***

([Aa]|[Bb]|[Cc])?-?\d-?\d?-?(O|N|alpha|beta|α|**β**)-? (linked)?

***Glycan Chain Type Descriptors***

Structural descriptors denoting glycan backbone or precursor chain types, often associated with histo-blood group antigens.

(Type-(1|2|3|4|I|II|III|IV)) (chain) (A|B|O|H|P|Pk|P1)

***Glycan Series Indicators (Prefix)***

neo | lyso | iso

(ganglio | globo | gluco | galacto | (iso)?globo | lacto)-?series

globo-?H

(cis | trans)-?

***Core Structure Descriptors***

Canonical glycan core structures and shorthand core notations.

core-?(1|2|3|4)-?

C1G

core-?fucos(ylated)?

***Antigenic Determinants (Prefix)***

Abbreviated forms of glycan antigens

.

(A | B | H | P | P1 | Pk)-?

***Glycan Compositional descriptors***

Structural classes describing glycan compositions and shorthand descriptors.

high-?mannose

hybrid

complex-?type

(man)\d

***Modification Patterns***

Terms describing enzymatic or chemical modifications of glycan structures, including residue addition, substitution, sulfation, phosphorylation, acetylation, etc.

sulfated | sulphated | sialylated | fucosylated | sialosylated | acetylated | galactosylated | pentaosylated | phosphorylated | core fucosylated

#### Class B: Named Glycan Motifs

Established glycan-related motifs such as blood group systems, canonical mentions of different antigens, and glycosaminoglycans (GAGs) are part of this class.

***Glycan-Related Blood Group Systems***

Standardized blood-group system names and related histo-blood group antigens are widely used in glycobiology.

(histo)?-?blood group(s)? (A | B | O | H | P | Pk | P1 | ABO | ABH)

(A | B | O | H | P | Pk | P1 | ABO | ABH) (histo)?-?blood group(s)?

***Lewis antigens***

Lewis antigens and a few standardized modified Lewis antigens are part of this.

(S|Sai|SD)?-?Lewis[^-\s]?\(?(a|A|b|B|x|X|y|Y)\)?(I|II|III|IV)?

(S|Sai|SD)?-?Le[^-\s]?\(?(a|A|b|B|x|X|y|Y)\)?(I|II|III|IV)?

SLX

SSEA-[1|2|3|4]

***Named glycan antigens and epitopes***

Established glycan antigens, carbohydrate epitopes, and glycan markers are used in glycobiology.

(T | Tn | TF | P | P1 | A | B | H | Pk | i | I | p^k | DF3 | CA19-9 | CA-50 | alpha-Gal | Forssman) (Ag | antigens)?

HNK-?(1|2|3) (antigens)?

STN-?(1|2|3) (antigens)?

Forssman glycolipid(s)?

CD10 | CD15 | CD26 | CD29 | CD44 | CD73| CD90 | CD105 | CD166 | CD174

***Named Glycosaminoglycans***

Named glycosaminoglycans and related carbohydrate polymers commonly occurring in glycoconjugates.

(chondroitin|dermatan|keratin|keratan|keraton|enoxaparin|heparin|heparan|bemiparin|bemiparan|certoparin|certoparan|parnaparin|parnaparan|reviparin|reviparan|sandoparin|sandoparan) (poly|mono)? (sulfate)? (A|B)?

(chondroitin|dermatan|keratin|keratan|keraton|enoxaparin|heparin|heparan|bemiparin|bemiparan|certoparin|certoparan|parnaparin|parnaparan|reviparin|reviparan|sandoparin|sandoparan)ic acid(s)

CS

DS

cytolipin (K|R)

hyaluronan acid(s)

hyaluronans

HA

fluxum

dermatan (L)? iduronate(s)

Cy222

**Class C: Glycobiology-derived Headwords**

General glycobiology-derived structural and biochemical headwords that are used to denote glycan components, glycoconjugate classes, antigenic features, and structural organization. These headwords can be combined with other classes to indicate a specific type of GST. Besides, they can be mentioned as individual terms in the literature.

alditol(s)? | determinant(s)? | core(s)? | chain(s)? | epitope(s)? | antigen(s)? | Ag | structure(s)? | saccharide(s)? | (N|O)?-?glycans | carbohydrate(s)? | cerebroside(s)? | ganglioside(s)? | cer(s)? | ceramide(s)? | globoside(s)? | globoseries | glycolipid(s)? | (glyco)?sphingolipid(s)? | sphingoshine(s)? | type(s)? | hapten(s)? | derivative(s)?

**Class D: Chemical Descriptors**

Chemical compounds that modify glycan structures.

***Chemical Substituent***

(ac | as | su | bis)-?

sialo | asialo | sulfo | sulpho

(sulfate|sulphate|phosphate)s?

HSO(3|4)

SO(3|4)

NH4

OH(4)?

CO(O)?

COOH

***Adjectival forms of chemical moieties***

Chemical moieties can be mentioned as adjectival forms to represent the modification of a GST.

(acetyl | phosphoryl |methyl) (I|II|III|IV)?

#### Class E: List of Sugar Names

Sugars are fundamental components of GSTs and play a central role in glycan composition. Since many GSTs are derived from sugar-related features, simple and modified sugar names, along with adjectival and aminyl-derived forms, are grouped within a unified Sugar class for consistent structural annotation and terminology interpretation.

***Monosaccharide List:***

Monosaccharides are the simplest form of carbohydrate, often called a "simple sugar." It cannot be broken down into smaller carbohydrate units through hydrolysis.

glyceraldehyde|dihydroxyacetone

(aldo|keto)?(tri|tetr|pent|hex|hept|oct|non)?ose(s)?

(erythr|thre|erythrul|rib|arabin|xyl|lyx|ribul|xylul|gluc|galact|mann|all|altr|gul|id|tal|fruct|sorb|tagat|pscic|sedoheptul|mannohelptul|glucohept|fuc|rhamn|quinov|digitox|abequ|tyvel|parat)ose(s)?

Gro|Dha|Ery|Thr|Eru|Rib|Ara|Xyl|Lyx|Rul|Xul|All|Alt|Glc|Man|Gul|Ido|Gal|Tal|Psi|Fru|Sor|Tag|Fuc|Rha|Sed|Mhu|Ghe|Qui|Dig|Abe|Tyv|Par

***Root forms of the Monosaccharides:***

tri|tetr|pent|hex|hept|oct|non|erythros|threos|erythrul|rib|arabin|xyl|lyx|ribul|xylul|gluc|galact|mann|all|altr|gul|id|tal|fruct|sorb|tagat|pscic|sedoheptul|mannohelptul|glucohept|fuc|rhamn|quinov|digitox|abequ|tyvel|parat

***Adjectival forms of the Monosaccharides:***

Root form + osyl

sialyl

***Amine forms of the monosaccharides:***

Root form + amine

GlcN|GalN|ManN|AllN|AltN|GulN|IdoN|TalN

GlcNAc|GalNAc|ManNAc|AllNAc|AltNAc|GulNAc|IdoNAc|TalNAc

***Aminyl forms of the Monosaccharides:***

Root form + aminyl

***Ketose/Aldose-related Monosaccharides:***

Root form + ulose

***List of Uronic Acids:***

Root form + uronic acid(s)

GlcA|GalA|ManA|IdoA|GulA|AltA|AllA|TalA

***List of Aldonic Acids:***

Root form + onic acid(s)

***List of Aldaric Acids:***

Root form + aric acid(s)

***List of Sialic Acids/Sialic Acid-like sugars:***

(poly)?sialic acid(s)

PSA

(poly)Sia

neuraminic acid(s)

pseudaminic acid(s)

legionaminic acid(s)

(Neu|NeuAc|NeuGc|Neu5Ac|Neu5Gc)-?H?-?\d?

Pse|Leg

KDO|KDN

***Other Sugar-like Acids:***

nonulosonic acid(s)

muramic acid(s)

MurNAc

***Monosaccharide Modifiers as Prefixes:***

*These* can be found before the root form of monosaccharides. Therefore, we also capture these prefix patterns so that the longer monosaccharide substring including the prefix can be detected.

(aldo|keto)?-?(D|L|alpha|beta|α|β|UDP|GDP|CMP|ADP|CDP|TDP|dTDP|deoxy|dideoxy)?-?

N?\,?N\'?-?(di)?(acetyl)?-?D?-?

N-?(glycolyl)?-?D?-?

***Disaccharide List:***

Disaccharides are made of two monosaccharide units linked together by a glycosidic bond.

sucrose|lactose|maltose|cellobiose|isomaltose|trehalose|melibiose|gentiobiose|laminaribiose|turanose|maltulose|palatinose|kojibiose|nigerose|sophorose|xylobiose|mannobiose|galactobiose|lactulose|leucrose|chitobiose|primeverose|vicianose|rutinose|neohesperidose|robinobiose|sambubiose|agarobiose|carrabiose|inulobiose|secalose|gentianose|erlose|theanderose|isomaltulose|trehalulose

Suc|Lac|Mal|Cel|IMal|Tre|Mel|Gen|Lam|Tur|Mltu|Pal|Koj|Nig|Soph|Xylb|Manb|Galb|Ltu|Leu|Chi|Pri|Vic|Rut|Neo|Rob|Sam|Aga|Car|Inu|Sec|Gnt|Erl|The|IMtu|Tru

***Root forms of the Disaccharides:***

sucr|lact|malt|cellobi|isomalt|trehal|melibi|gentiobi|laminaribi|turan|maltul|palatin|kojibi|niger|sophor|xylobi|mannobi|galactobi|lactul|leucr|chitobi|primever|vician|rutin|neohesperid|robinobi|sambubi|agarobi|carrabi|inulobi|secal|gentian|erl|theander|isomaltul|trehalul

***Other Saccharides:***

cellotriose|raffinose|panose|kojitriose

stachyose

verbascose

***Disaccharide motifs:***

lactosamine(s)?

lactosediamine(s)?

lacto-N-biose

LacNAc|LacdiNAc

(Lac|Lacdi)-?NAc-?H?-?\d?

LN(T)?-?\d?

(LDN|LDN-F)-?(1|2)?

(Neu5Ac[αa]2-\d)?Gal(NAc)?[βb]1-\d(\(Fuc[αa]1-\d\))?GlcNAc

#### Class F: Glycoconjugates

A glycoconjugate (GCT) is a molecule in which a sugar is linked to another chemical group (aglycon) through a glycosidic bond. This class contains all the named GCTs used in glycan terms, as well as subclasses of GCTs such as glycosphingolipids (GSLs), globosides, etc., where the number of sugar molecules varies along with the chemical group.

***Named Glycosides***

These are specific glycosides available in glycobiology. Other sub-classes of glycosides, such as GSLs are part of it. Each of these glycosides has several other child classes.

(gangli | lact | glyc | mann | fuc | cerebr | sial | hex | fruct | xyl | all | altr | gul | id | tal | arabin | rib | lyx | rhamn | digitox)oside(s)?

***Glycosphingolipids***

Glycosphingolipids (GSLs) are ceramides with attached glycan structures. Some of the glycosides are known as GSLs (e.g., globoside, cerebroside, ganglioside). GSLs are also categorized into several subclasses. For example:

***i) Named Gangliosides***

GSLs that contain sialic acids are part of this subclass.

(GM|GT|GQ|GD|GA)-?\d(a|b|c)?(a|b|c)?

(LD|LE|LQ|TD|TE)-?[1|2](a|b)?

TQ-?[1](a|b)?

CD[1|2][1|2]b

***ii) Named Cerebroside***

A single sugar attached to the ceramides is part of it. Adjectival forms of simple sugars ending with ceramides are part of it.

(glycosyl|glucosyl|galactosyl|mannosyl|fucosyl)ceramide

ceramide monohexoside(s)?

CMH

***iii) Neutral Oligosaccharide Glycosphingolipids, Globosides and Glycolipids***

Any neutral glycans attached to the ceramides are part of this subclass. Besides GSLs, globosides and glycolipids are also part of this type.

ceramide (tri|di|tetra)hexoside

CTH | CDH | CTeH | CPH

(globo)?(neo)?(lactos | triaos | trios | tetraos | hexos | hexaos | pentaos | heptos | heptaos | octaos)yl[-\s]?ceramide(s)?

n?Lc[-_]?(3|4|5)?

(DS)?i?Gb\d?

GL[-_]?\d?

(iso)?globoside(s)? (1|2|3|4|I|II|III|IV))?

***iv) Sulfated Glycosphingolipids***

This is a special class that contains sulfated glycosphingolipids.

Sulfatide(s)?

#### Class G: Non Specific

cyclic | para | pauci | Sd(A) | Sd^A | Cad | seminolipid

VIM-[1|2]
